# Phylophenomic and Phylogenomic analysis for *Ovis aries* reveals distinct identity of newly reported breed

**DOI:** 10.1101/2022.07.31.502249

**Authors:** Aruna Pal, Samiddha Banerjee, Prabir karmakar

## Abstract

Domestication and phylogenetics for Ovis aries is an important species to study, since there exists enormous biodiversity in terms of habitat and utility of sheep. The present study aimed at identification of the biodiversity existing within sheep breeds reared in different agroclimatic zones of the state West Bengal (Garole, Birbhum, Bonpala and Chotanagpuri) through phylogenetic analysis of phenotypic traits as growth and biomorphometric traits through principal component analysis, factor analysis, genetic correlation, multivariate cluster analysis through Hierarchial classification and k-means cluster analysis. Confirmation of the phylophenomic studies were later on carried out with phylogenomic analysis with microsatellite markers for sheep. Birbhum sheep from dry arid region of West Bengal is reported as the newly reported breed of sheep with distinct genetic identity.

## 1. Introduction

It is quite interesting to study the domestication and evolution leading to biodiversity in Ovis aries, since it has been reported that sheep and dog were the first animal to be domesticated^**1**^. It has been observed that sheep are distributed across the globe and are reared in both tropical and temperate region. They are available in every agroclimatic conditions, hilly region, terai, alluvial, dry arid and coastal region. Moreover, another important and interesting feature is that sheep are reared for varied purposes of economic importance. In some places, they are reared for wool (high quality apparel wool or carpet wool), while in other places, mutton production is important. Even in some places, they are reared for milk. Keeping the above points, it is quite evident that biodiversity among sheep exists and it is quite important to study the phylogeny and evolution of sheep.

In our lab, we had reported the phylogenetic analysis for sheep initially with mitochondrial gene as Cytochrome B^**2**^. Later on, phylogenomic analysis for sheep were conducted with complete mitochondrial genome^**3**^. Indigenous breeds of sheep are mostly adapted to their local climatic conditions and somewhat resistant to diseases. They have tremendous socioeconomic importance in rural economy for marginal and landless farmers. Genetic characterization through immune response genes and other parameters have been attempted in our earlier studies^**2-12**^. It has been widely used in studies of breed characterization and genetic diversity as it provides a descriptive analysis of the differences between populations, considering all variables together, providing a data overview^**13-14**^. From our lab, we had already studied growth parameters or biomorphometric characteristics for other ruminant as cattle & buffalo^**15-17**^, goat^**2-18**^ and documented duck^**19**^. A thorough study on one of the important gene responsible for growth as growth hormone gene has been conducted and certain interactions with growth and other phenotypic traits have been revealed^**16-17, 20-23**^.

Phylophenomic studies refer to the phylogenetic analysis with phenotypic traits. Growth traits along with biomorphometric characteristics are the promising traits for such analysis through multivariate data analysis. Multivariate Data Analysis refers to a statistical technique used to analyze data from more than one variable. In case of quantitative inheritance of livestock, the traits are governed by polygenic inheritance and hence multiple data are of prime importance. Despite the quantum of data available, the ability to obtain a clear picture of what is going on and make intelligent decisions is a challenge. When available information is stored in database tables containing rows and columns, Multivariate Analysis can be used to process the information in a meaningful fashion^**24-25**^. Cluster analysis is a data exploration (mining) tool for dividing a multivariate dataset into “natural” clusters (groups). The method is useful to explore whether previously undefined clusters (groups) may exist in the dataset and is used when the sample units come from an unknown number of distinct populations or sub-populations with no apriori definition of those populations. In the present study, we hypothesize that different breeds within a species will be clustered separately since the individuals within a breed are genetically more similar compared to individuals between two different breeds.

In a similar way, phylogenomic analysis with microsatellite markers are of importance. Microsatellites are the short segment of DNA (one to six or more base pairs in length) which is repeated in succession at a particular genomic location. These DNA sequences correspond to the non-coding region and are generally designated as junk DNA.

Keeping the above facts in view the present study was aimed at characterization of the sheep population of West Bengal and biodiversity study through multivariate data analysis and microsatellite genotyping of sheep reared under the different agroclimatic zone of West Bengal.

## 2. Materials and Method

### Animals and area of study

The present study covered randomly chosen 2254 sheep maintained by marginal farmers as well as Government Farms in different agroclimatic zones of West Bengal (Table 1). The data represents adult sheep (12 ± 2 m of age) for growth traits.

**Table 1.** Agroclimatic variables and study area.

| Agroclimatic regions | Blocks covered in study |  | Latitude (N) | Longitude (E) | Avg. temp. (°C) |
| --- | --- | --- | --- | --- | --- |
| Terai | Jalpaiguri | Ramsai Farm<br>Rajganj | 26°15'47" to<br>25°55'34" | 88°23'22" to<br>89°7'30" | 7.8 to<br>37.9 |
| Old-Alluvial | Dinajpur<br>(S) | Balurghat | 26°35'15" to<br>25°10'55" | 85°00'30" to<br>87°48'37" | 20.4 to<br>31.4 |
| New-Alluvial | Nadia | Haringhata<br>State Farm, Kalyani.<br>LFC Farm, WBUAFS | 22°52'30" to<br>24°05'40" | 22°08'10" to<br>88°48'15" | 20.6 to<br>32.4 |
| Red-Laterite | Birbhum<br>Bankura | Dubrajpur,<br>Khoyrasole, Rajnagar<br>Indpur | 23°32'30" to<br>24°35'00" | 88°1'40" to<br>87°5'25" | 15 to<br>40 |
| Coastal-Saline | 24-Pgs (S) | Sagar, Gosaba | 21°25'30" to<br>23°16'50" | 88°01'10" to<br>89°06'15" | 16-43 |

Recording of other sheep breeds of eastern India (i.e. Sahabadi, Ganjam, Balangir, Kendrapada and Tibetan sheep) were collected from their respective breeding tract for necessary comparison with existing populations of West Bengal.

### Growth and Biomorphometry traits

Growth and biomorphometric parameters recorded includes body weight (BW) and daily body weight gain (DBWG), heart girth (HG), body length (BL), body height (BH), pelvic width (PW), tail length (TL), head length (HL), ear length (EL), ear width (EW), distance between two eyes as defined by FAO, 2012^**26**^. BW was measured at site with spring balance whereas other biomorphometric characters were measured with measuring tape.

### Statistical Analysis

Mean and standard error of the traits were calculated using standard statistical procedure^**27**^. For the mean comparison of each biomorphometrical characteristic for different populations one way analysis of variance was used and the pairwise comparison of means was done by using Duncan Multiple Range Test^**28**^.

### Phenotypic correlation

The phenotypic correlations (r_p_) between records of two traits was estimated by using the following formula:

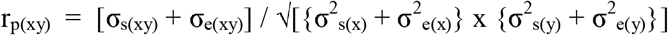

The standard errors of phenotypic correlation were obtained by using the following formula:

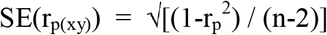

The significance of phenotypic correlations was adjudged after estimating the t-values and comparing with table values at (n-2) d.f. as given by Snedecor and Cochran (1967)^**27**^.

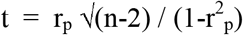

Pearsons correlation was estimated through SYSTAT package.

### Principal component analyses

Principal component analysis is used when the recorded traits are correlated. Principal component analyses (PCA) is a multivariate approach, which reduces the dimensionality of a data set. The PCA can explain relationships between different body biomorphometry traits in a better way. SPSS 17 software was used for the analysis. PCA is a multivariate statistical techniques dealing with the reduction of a set of observable variables. PCA accounts for the maximum portion of the variance present in the original set of variables with a minimum number of composite variables. Varimax rotation was employed for rotation of principal components. The transformation of the components were employed to approximate a simple structure. Kaiser-Meyer-Olkin (KMO) test of sampling adequacy^**29**^ and Bartlett’s test of sphericity were computed to establish the validity of data set, at 1% level of significance.

### Factor analysis

Factor analysis is a statistical technique used for data reduction. It is used to describe variability among observed, correlated variables in terms of a potentially lower number of unobserved variables called factors. Factor analysis assumes that a variable’s variance can be decomposed into two parts^**30**^. These are common variance (communality factor) that is shared by other variables included in the model; and specific variance (unique factor) as it is specific to a particular variable and includes the error variance. The estimate of communality for each variable measures the proportion of variance of that variable explained by all the other components jointly. It assumes that the unique variance contributes a small portion of the total variance^**31**^. Scree plot was obtained from Systat13 software.

### Cluster analysis by Hierarchical Classification

Multivariate Cluster Analysis was employed for estimation of genetic distance and construction of phylogenetic tree for sheep population at different agroclimatic regions of West Bengal with reference to the native population of eastern India (SYSTAT 13) using the traits under study. Genetic distance studies and phylogenetic tree was constructed based on Euclidean distance and Complete Linkage Method^32^. Euclidean distance analysis is one of the multi-variate analysis by which a population can be classified into subgroups (clusters) using various traits, so that the sheep in a particular cluster are similar, but different from the animals of other clusters. Hierarchical classification was employed for phylogenetic analysis among different sheep breeds of eastern India.

The genetic distance was calculated by the following method as suggested by Jobson (1992)^**32**^

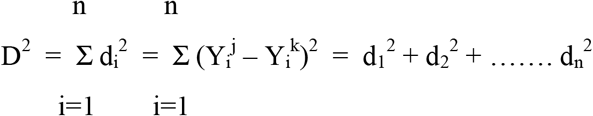

| Genotype | Y<sub>1</sub> | Y<sub>2</sub> | .... | Y<sub>n</sub> |
| --- | --- | --- | --- | --- |
| Bulls genotype (J) | .... | .... | .... | .... |
| Bull's genotype (K) | .... | .... | .... | .... |
|  | d<sub>1</sub> | d<sub>2</sub> | .... | d<sub>n</sub> |

Where,

d^2^ = Squared genetic distance between two genotypes,

Y_i_ = ith character of the sheep,

j = One genotype of the gene, and

k = Another genotype j ≠ k.

### K means cluster analysis

K-**means clusteranalysis** partition n observations into **k clusters** resulting in a partitioning of the data space into Voronoi cells. K means clustering was also employed for differentiating Birbhum and Garole sheep based on biomorphometry of traits in software package of Systat 13.

K-means is one of the simplest unsupervised learning algorithms that solve the well known clustering problem^**33**^. The basic concept is to define k centroids, one for each cluster. The next step is to take each point belonging to a given data set and associate it to the nearest centroid. When no point is pending, the first step is completed and an early groupage is done. At this point we need to re-calculate k new centroids as barycenters of the clusters resulting from the previous step. After we have these k new centroids, a new binding has to be done between the same data set points and the nearest new centroid. A loop has been generated. As a result of this loop we may notice that the k centroids change their location step by step until no more changes are done. In other words centroids do not move any more. Finally, this algorithm aims at minimizing an *objective function*, in this case a squared error function. The objective function

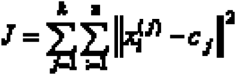

Where 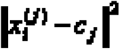 is a chosen distance measure between a data point 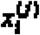 and the cluster centre *c*_*j*_, is an indicator of the distance of the *n* data points from their respective cluster centres.

### Microsatellite genotyping for Ovies aries

Blood samples were collected from 40 unrelated animals (rams and ewes) of Birbhum sheep in its breeding tract. The genomic DNA from the blood samples was isolated using a standard phenol/chloroform/isoamyl alcohol extraction method (Sambrook *et al*. 1989).

PCR amplification set of 24 microsatellite markers based on the list of MoDAD (FAO)^**34**^ were utilized to generate data on DNA samples of the Birbhum sheep. The details of microsatellite markers, primer sequences, size range, gene bank accession number and chromosomal location is given in Table 4. The forward primer for each marker was fluorescently labeled with FAM, NED, VIC or PET dye. Amplification of the loci (multiplexed) was performed in a 25µl final reacon volume containing at least 100 ng of genomic DNA, 5 pico moles / µl of each primer, 1.5mM MgCl, 200µM dNTPs, 0.5 U Taq DNA polymerase and 1x Taq buffer. A common touch 2 down PCR programme as suggested under MoDAD project^**26**^ without extension step was used for the amplification of all the twenty four markers. PCR amplification consisted of 3 cycles of 45 sec at 95 C, 1 min at 60 C; 3 cycles of 45 sec at 95 C, 1 min at 57 C; 3 cycles of 45 sec at 95 C, 1 min at 54 C; 3 cycles of 45 sec at 95 C, 01 min at 51 C and 20 cycles of 45 sec at 95 C, 1 min at 48 C. The amplified products were resolved on 2% agarose gel.

### Genotyping of the PCR products

PCR products were genotyped on an automated DNA sequencer using LIZ 500 as internal lane standard (ABI PRISM). Gene Mapper software version 3 was used to extract raw data. Popgen 3.2^**35**^) and GenAlEx6.5^**36**^ softwares were used for the genetic diversity analysis. Polymorphism Information Content (PIC) of the microsatellite loci was estimated^**37,38**^. The primers used for microsatellite genotyping for sheep are depicted in Table 2.

**Table 2:**
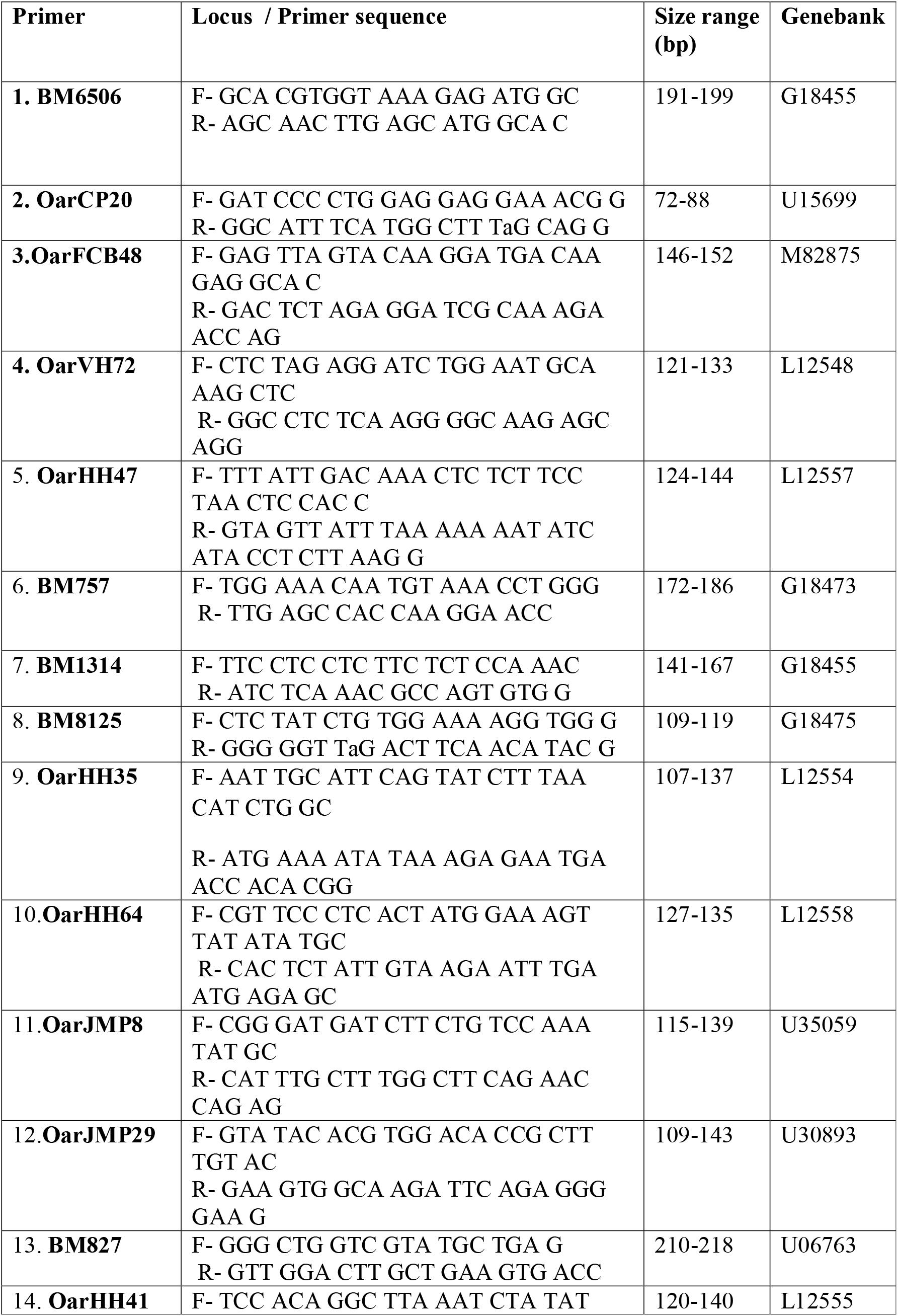
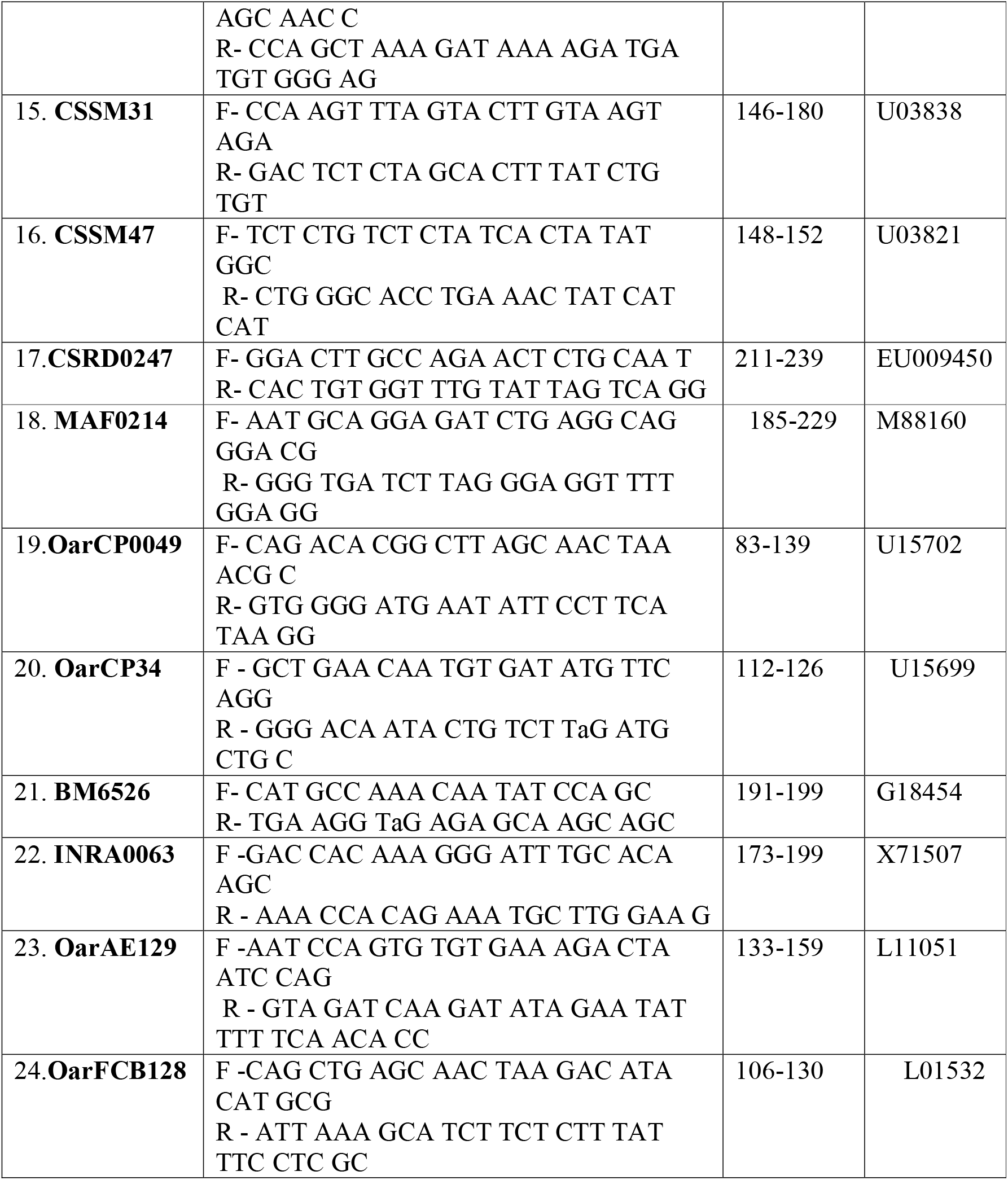
Microsatellite marker for sheep as per FAO.

## 3. Result

### Growth traits of sheep reared at varied agroclimatic region

Significant difference among the growth traits for different sheep breeds of West Bengal has been observed and characterized accordingly (Table 3). The average growth and biomorphometric characters from different agroclimatic zone for sheep breed, ram (Fig 1) and ewe (Fig 2) have been depicted. A new breed of sheep was identified as Birbhum from Birbhum district of West Bengal, breeding tract as depicted in Fig 3a and photographs of ram, ewe and Birbhum sheep with rudimentary ear have been presented in Fig 3b. Since Garole and Birbhum seem to be genetically closer, a detailed study has been undertaken for these two breeds. Several biometric traits found to differ significantly which includes BW, HG, PG and BH. Birbhum sheep under the present study exhibited significantly better BW and BH compared to the Garole but HG and PG remained superior for the Garole, because of the compactness of body structure of Garole. Similar trend was observed for either sex within the breed under study. Male always excelled female in most of the growth traits, but female had better ear length and ear width in both Garole and Birbhum sheep. In the present study 30.6 % of the Birbhum sheep found to have rudimentary ear, which may be an unique characteristics for this particular breed. Body colour varied from black, brown and white for the Birbhum sheep. Horn is present in ram but absent in ewe in both Garole and Birbhum breed. Average ear length and ear width was exceptionally less for Birbhum in comparison to Garole might be due to higher percentage of rudimentary ear. Hence Garole and Birbhum sheep have distinctly separate morphological characteristics.

**Table 3:** Growth parameters for different sheep breeds of West Bengal.

|  | Garole | Bonpala | Chotanagpuri | Birbhum | P value |
| --- | --- | --- | --- | --- | --- |
| Body weight (kg) | 10.43±0.39<sup>a</sup> | 41.71± 0.32<sup>c</sup> | 19.45± 0.19<sup>d</sup> | 12.97±0.33<sup>b</sup> | 0.946 |
| Average daily body weight gain (gm per day) | 101.039 ±2.86 | 90.86±2.96 | 95.43±3.56 | 115.53± 2.45 |  |
| Body length (cm) | 45.91±0.39<sup>a</sup> | 74.87±0.63<sup>b</sup> | 52.93±0.28<sup>c</sup> | 47.73 ± 0.29<sup>a</sup> | 0.62 |
| Heart girth (cm) | 62.97±0.42<sup>a</sup> | 80.73±0.74<sup>c</sup> | 71.19±0.62<sup>d</sup> | 60.57± 0.38<sup>b</sup> | 0.62 |
| Paunch girth (cm) | 63.74±0.41<sup>a</sup> | 81.93±0.69<sup>c</sup> | 72.67±0.57<sup>d</sup> | 60.86 ± 0.48<sup>b</sup> | 0.909 |
| Body height (cm) | 47.67±0.39<sup>a</sup> | 73.39±0.53<sup>c</sup> | 54.64±0.53<sup>d</sup> | 51.48± 0.37<sup>b</sup> | 0.831 |
| Head length (cm) | 15.61±0.39 | 18.99±0.43 | 18.07±0.44 | 17.58± 0.39 |  |
| Tail length (cm) | 10.01±0.38 | 15.41±0.39 | 13.20±0.23 | 11.78± 0.39 |  |
| Ear length (cm) | 6.49±0.37 | 7.92±0.23 | 9.40±0.24 | 4.88± 0.47 |  |
| Ear Width (cm) | 4.66±0.36 | 3.57±0.35 | 4.22±0.39 | 3.89± 0.30 |  |
| Distance between two eyes (cm) | 11.03±0.37 | 13.69±0.67 | 12.54±0.48 | 9.16± 0.39 |  |

\*Mean values bearing different superscripts (a,b,c,d) in a row differ significantly . The data represents the value of average of male and female

**Table no. 4:**
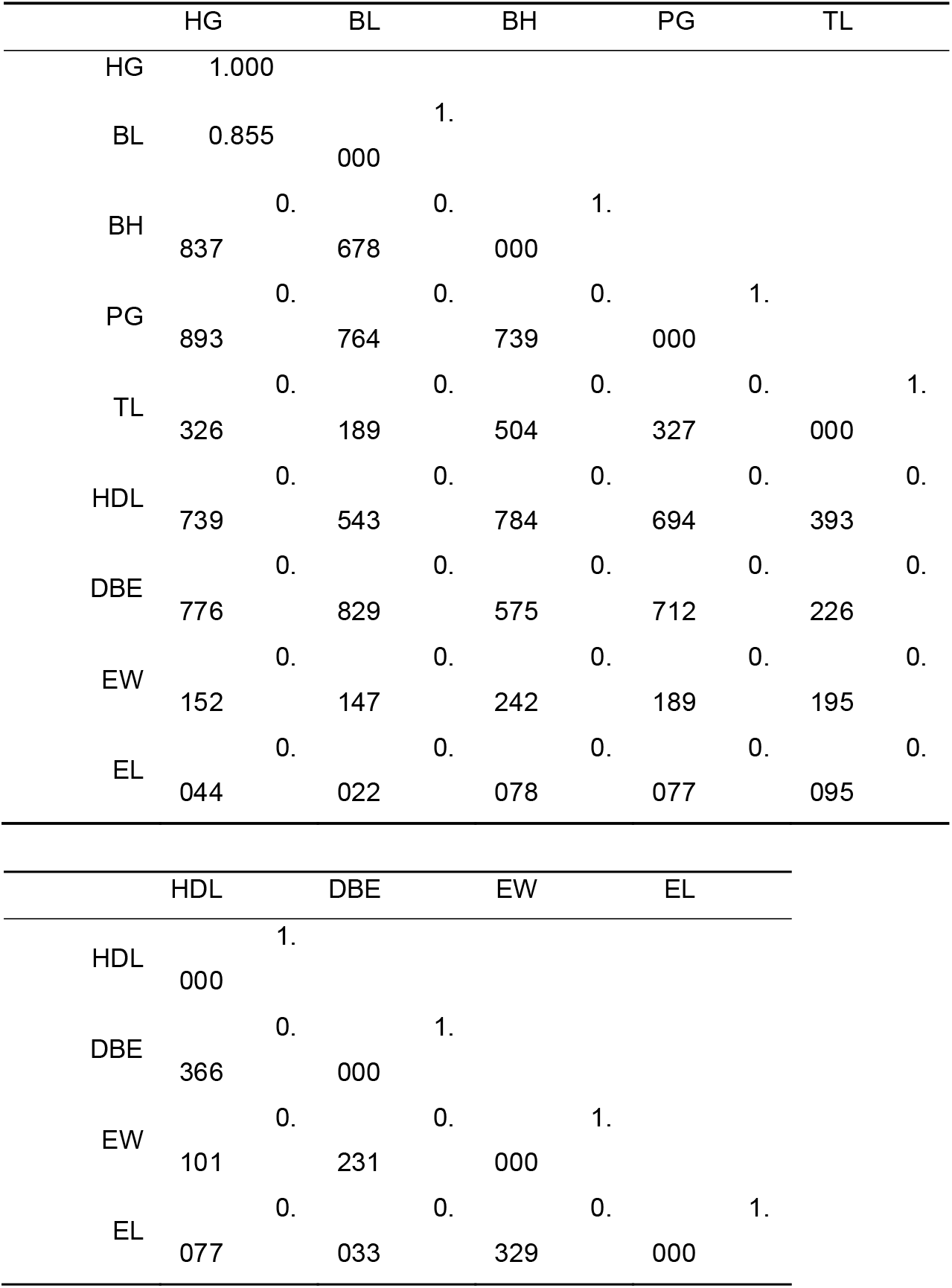
Phenotypic correlation between growth and morphometric traits at 12 months of age in sheep.

**Fig 1:**
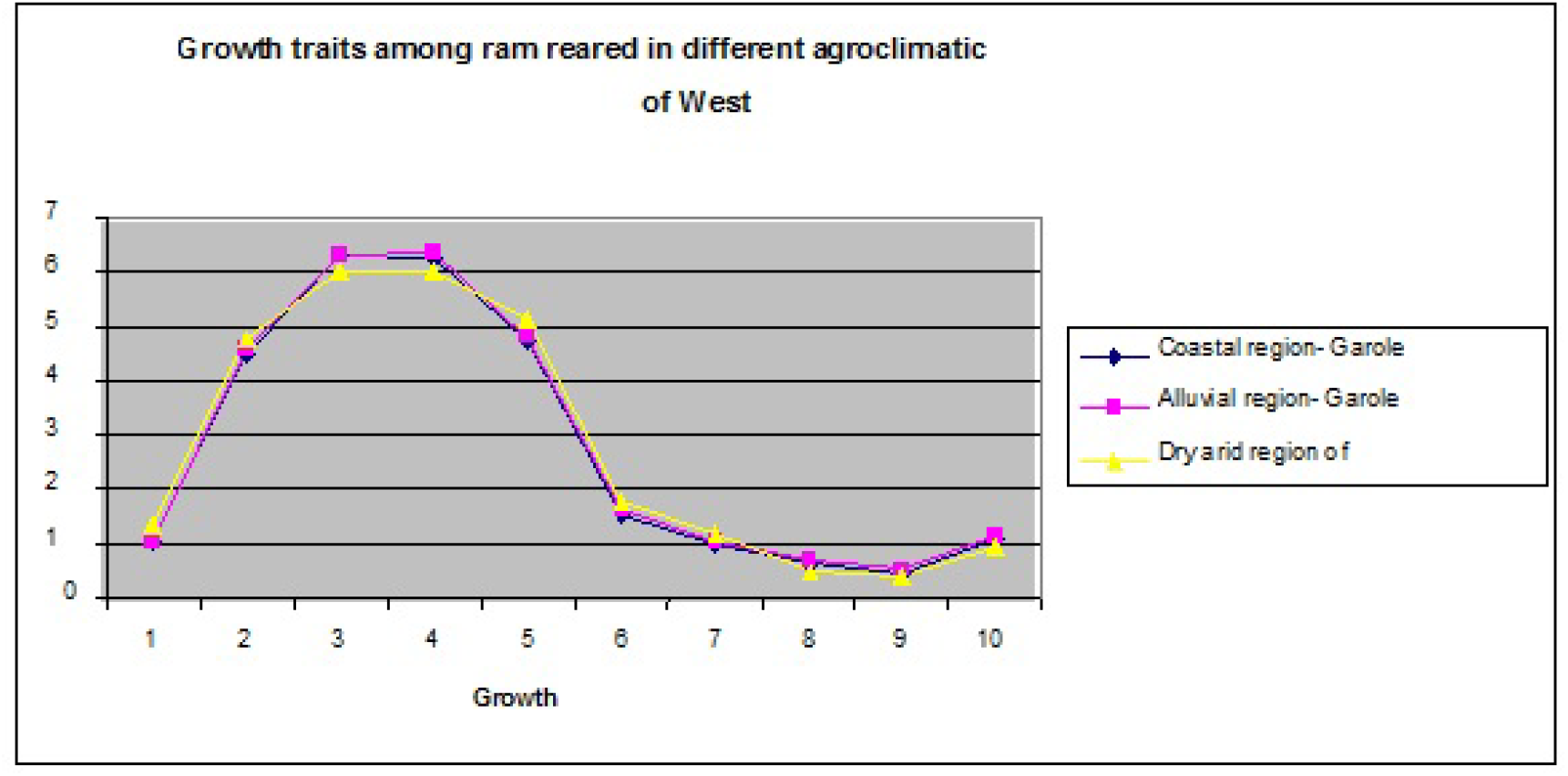
Growth traits among male sheep at different agroclimatic region of West Bengal 1. Body weight (kg) 2. Body length (cm) 3. Heart girth (cm) 4. Paunch girth(cm) 5. Body height(cm) 6. Head length(cm) 7. Tail length(cm) 8. Ear length(cm) 9. Ear Width(cm) 10.Distance between two eyes (cm)

**Fig 2:**
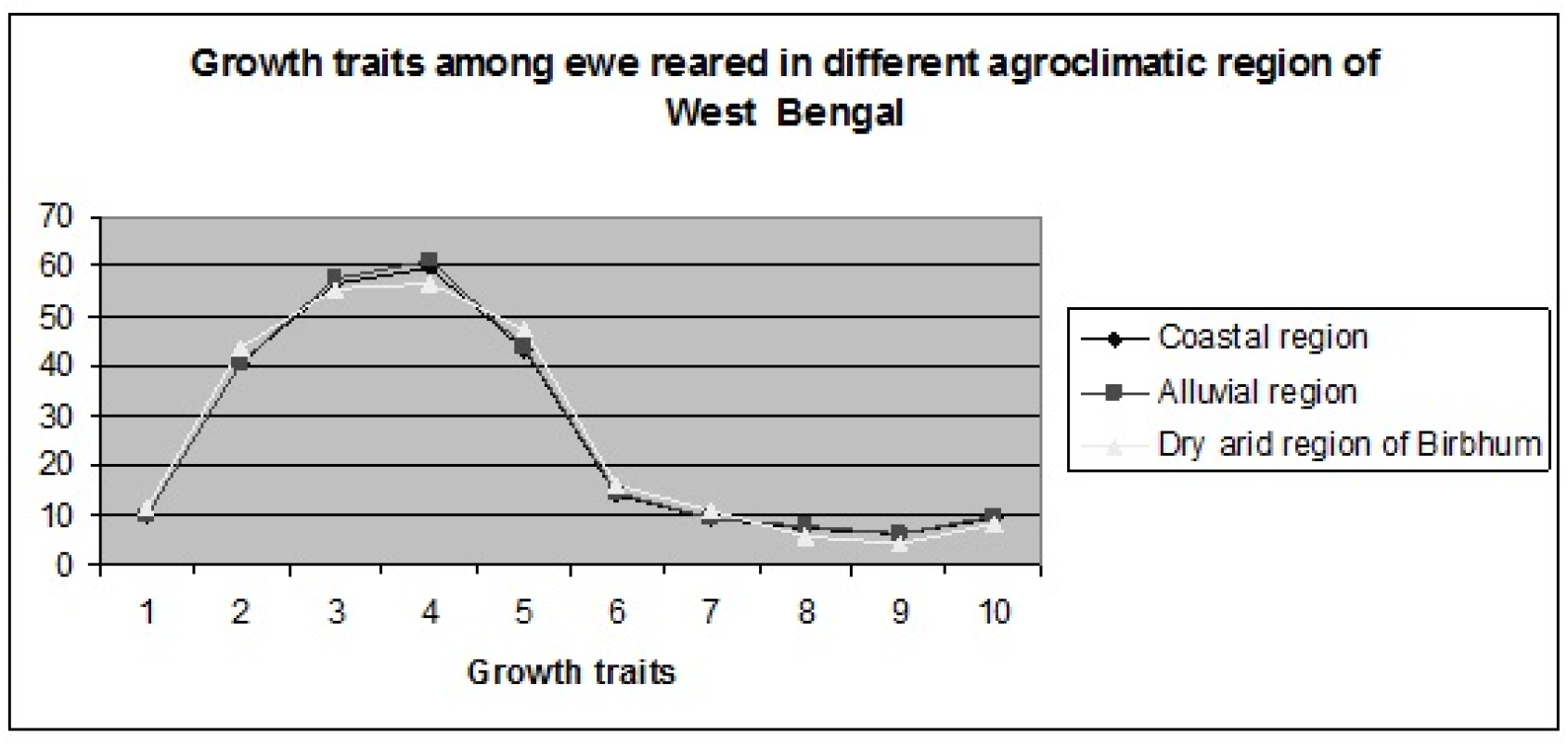
Growth traits among female sheep at different agroclimatic region of West Bengal 1. Body weight (kg) 2. Body length (cm) 3. Heart girth (cm) 4. Paunch girth(cm) 5. Body height(cm) 6. Head length(cm) 7. Tail length(cm) 8. Ear length(cm) 9. Ear Width(cm) 10.Distance between two eyes (cm)

**Figure 3a:**
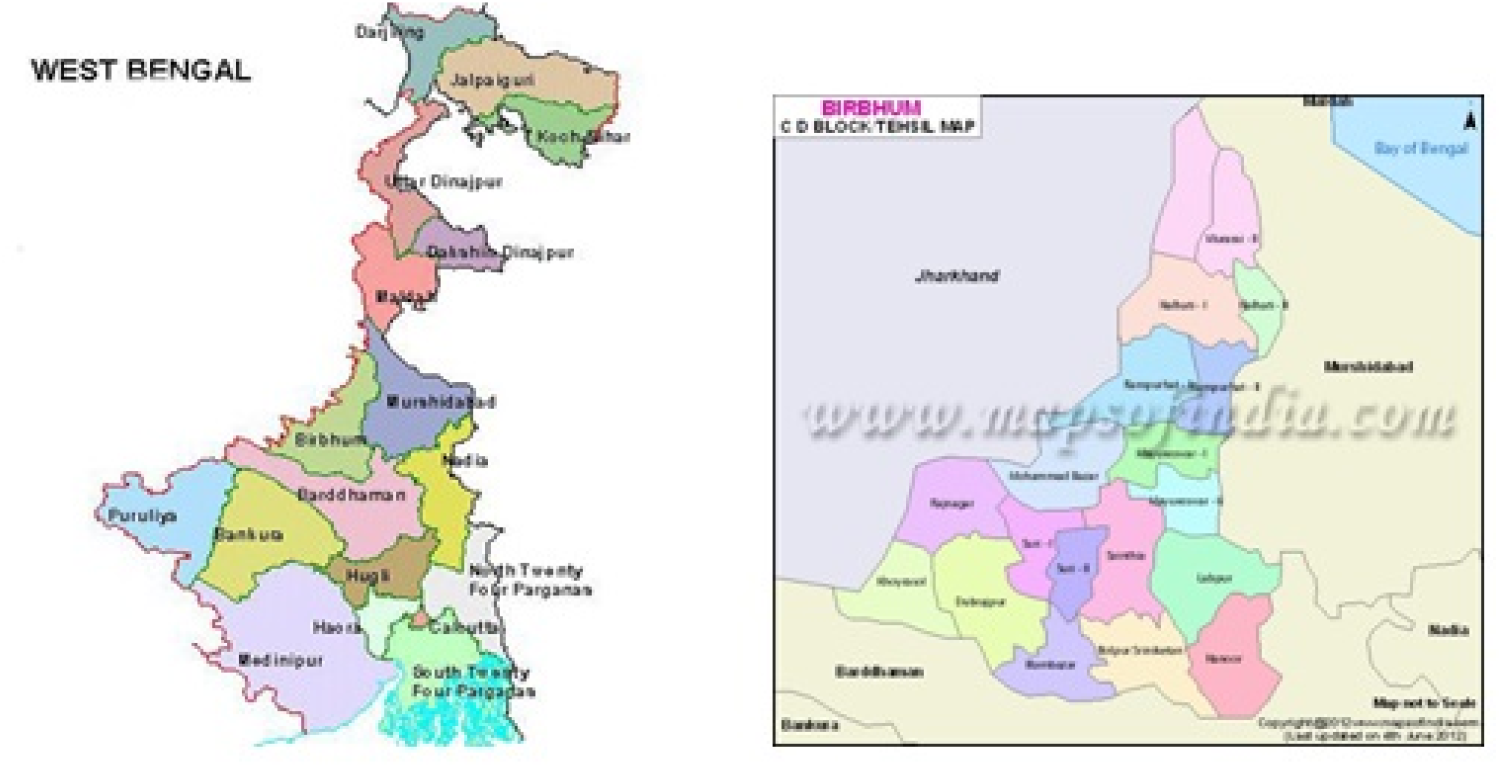
Map of West Bengal and that of Birbhum district

**Figure 3b:**
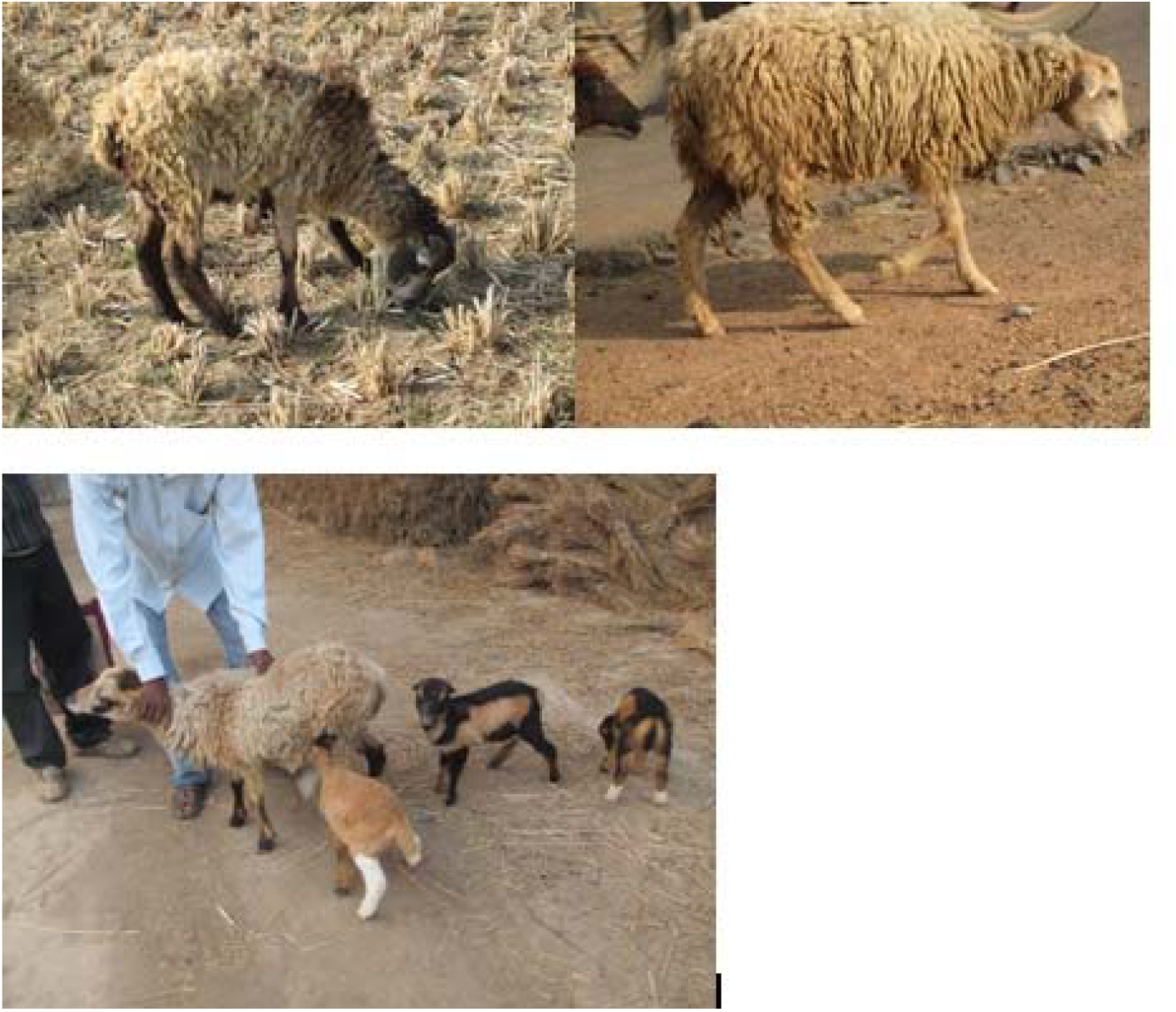
Photograph of Birbhum ram (left) & ewe (Right). Birbhum sheep with rudimentary ear (bottom), which is inherited to lambs.

*Mean values bearing different superscripts (a,b,c,d) in a row differ significantly. The data represents the value of average of male and female

Figure 3a depicts the map of West Bengal and that of Birbhum district & Figure 3b depicts Photograph of Birbhum ram (left) & ewe (Right). Birbhum sheep with rudimentary ear (bottom), which is inherited to lambs.

### Phenotypic correlation among the different growth parameters

Phenotypic correlation between morphometric traits of sheep at 12 month of age is presented in Table no. 4 as Pearson correlation co-efficient. Significant positive correlation was observed for most of the growth traits in sheep at 12 months of age under study. The graphical representation of phenotypic correlation has been presented in Fig 4.

**Figure 4:**
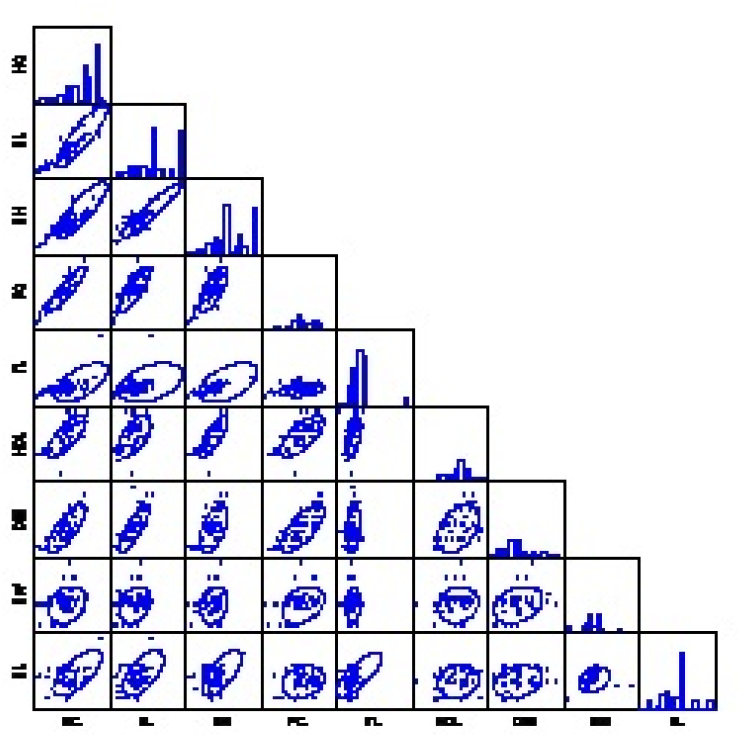
Graphical representation of correlation (Pearson’s correlation data) for biomorphometric traits of sheep

Significantly high and positive correlation was observed between body weight with body length, body height, heart girth, punch girth and head length. Correlation between heart girth and body weight was found to be significantly higher at 12 months of age.

### Principal component analysis

The PCA was applied to 10 body conformation traits in sheep population of West Bengal from different agroclimatic zone. Out of 10 principal components, two components were extracted using Kaiser Rule criterion to determine the number of components i.e. retaining only the components that have eigen value greater than 1 (Table 5). The Varimatrix rotated component matrix of different factors for biometric traits for sheep in PCA have been represented in Table 6. Scree plot is useful to decide the various components and the actual number of the components to be included for analysis. Components having eigenvalues up to the point “bent of elbow” are usually considered (Figure 5). The identified two components could explain cumulative percentage of variance of 84.67%. First component accounted for 66.67% of the variation. It was represented by significant positive high loading of heart girth, body height, body length, punch girth, head length, distance between eyes and body weight. First component seemed to be explaining the maximum variability among the sheep. So, the identified traits under first component were further utilized for hierarchical classification analysis.

**Table 5:** Total Variance Explained by different components of Principal component analysis.

| Component | Initial Eigenvalues |  |  | Extraction Sums of Squared Loadings |  |  |
| --- | --- | --- | --- | --- | --- | --- |
|  | Total | % of Variance | Cumulative % | Total | % of Variance | Cumulative % |
| 1 | 6.686 | 66.857 | 66.857 | 6.686 | 66.857 | 66.857 |
| 2 | 1.781 | 17.813 | 84.670 | 1.781 | 17.813 | 84.670 |
| 3 | .993 | 9.933 | 94.604 |  |  |  |
| 4 | .215 | 2.148 | 96.752 |  |  |  |
| 5 | .136 | 1.363 | 98.115 |  |  |  |
| 6 | .113 | 1.126 | 99.241 |  |  |  |
| 7 | .038 | .383 | 99.624 |  |  |  |
| 8 | .019 | .191 | 99.815 |  |  |  |
| 9 | .016 | .156 | 99.971 |  |  |  |
| 10 | .003 | .029 | 100.000 |  |  |  |

**Table 6:** Varimatrix rotated component matrix of different factors for biometric traits for sheep in PCA

|  | Component |  |
| --- | --- | --- |
|  | 1 | 2 |
| HG | .961 | -.023 |
| BH | .946 | -.110 |
| BL | .979 | -.129 |
| PG | .945 | -.033 |
| TL | .463 | .261 |
| HDL | .944 | -.025 |
| DBE | .921 | -.108 |
| EW | .450 | .868 |
| EL | .204 | .904 |
| BWT | .905 | -.318 |

**Figure 5:**
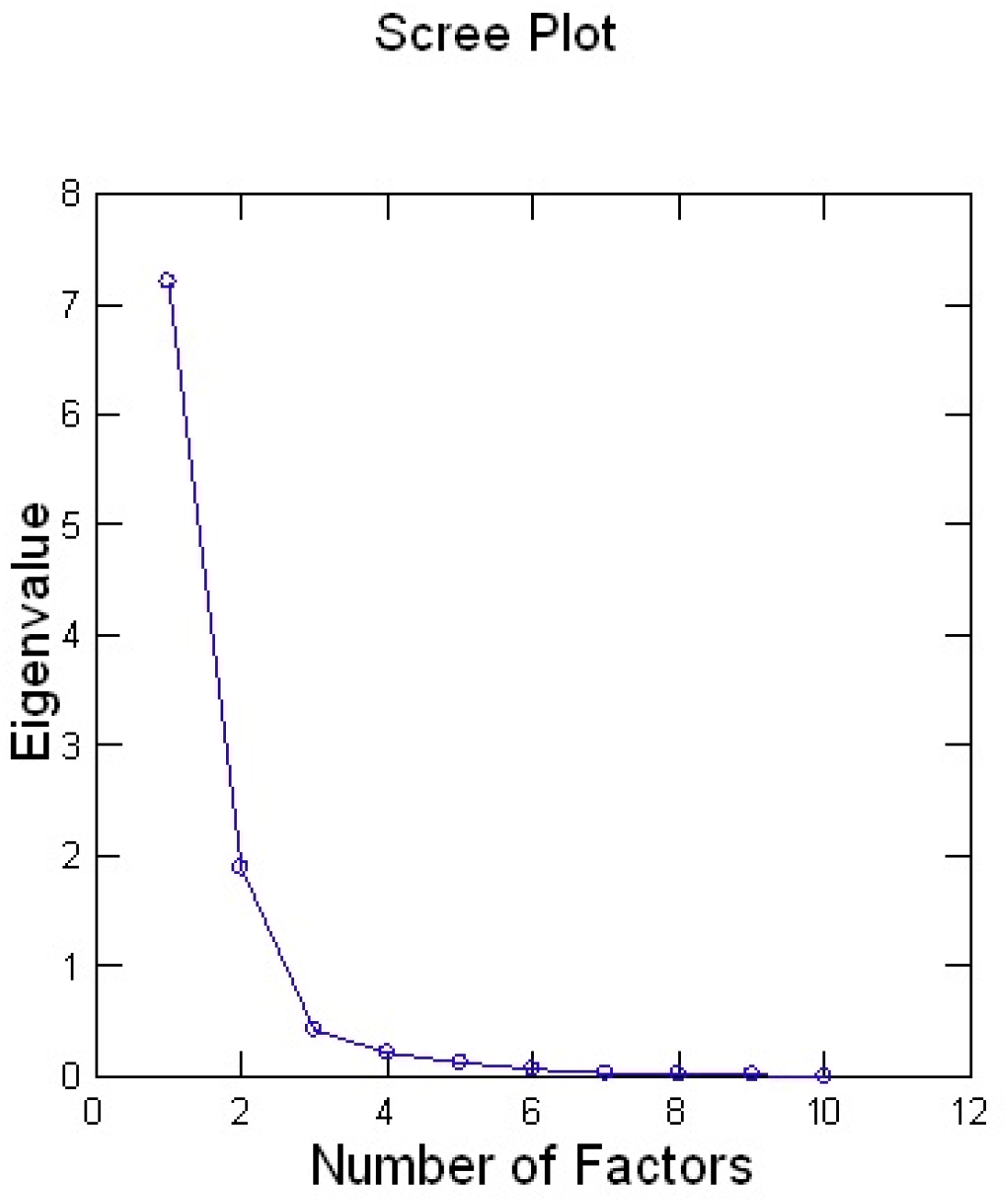
Scree plot showing component number with eigen values in Principal component analysis

### Factor analysis

The observation was similar to Principal component analysis. Only two components have been identified with eigen value greater than one. Component one explained 53.928 of the total variance (Table 7). Scree plot also depict the various components and could can be used to decide the actual number of the components to be included for analysis, components having eigenvalues up to the point “bent of elbow” are usually considered (Figure 6a). The identified two components could explain cumulative percentage of variance of 68.64%. First component accounted for 53.93% of the variation. It was represented by significant positive high loading of heart girth, body height, body length, punch girth, head length, distance between eyes and body weight. First component seemed to be explaining the maximum variability among the sheep. Factors loadings plot obtained from factor analysis have been graphically represented in Fig 6b. Factor analysis was used in addition to principal component analysis for further confirmed the result.

**Table 7:** Total variance explained by factor analysis.

**Latent Roots (Eigenvalues)**
|  | 1 | 2 | 3 | 4 | 5 |
| --- | --- | --- | --- | --- | --- |
|  | 4.853 | 1.324 | 1.000 | 0.716 | 0.540 |
|  | 6 | 7 | 8 | 9 |  |
|  | 0.223 | 0.155 | 0.130 | 0.059 |  |

**Component loadings**
|  | 1 | 2 |
| --- | --- | --- |
| HG | 0.960 | 0.138 |
| BL | 0.867 | 0.192 |
| BH | 0.894 | -0.047 |
| PG | 0.910 | 0.078 |
| TL | 0.461 | -0.314 |
| HDL | 0.785 | 0.019 |
| DBE | 0.800 | 0.109 |
| EW | 0.273 | -0.734 |
| EL | 0.109 | -0.781 |

**Variance Explained by Components**
|  | 1 | 2 |
| --- | --- | --- |
|  | 4.853 | 1.324 |

**Percent of Total Variance Explained**
|  | 1 | 2 |
| --- | --- | --- |
|  | 53.928 | 14.715 |

**Fig 6a:**
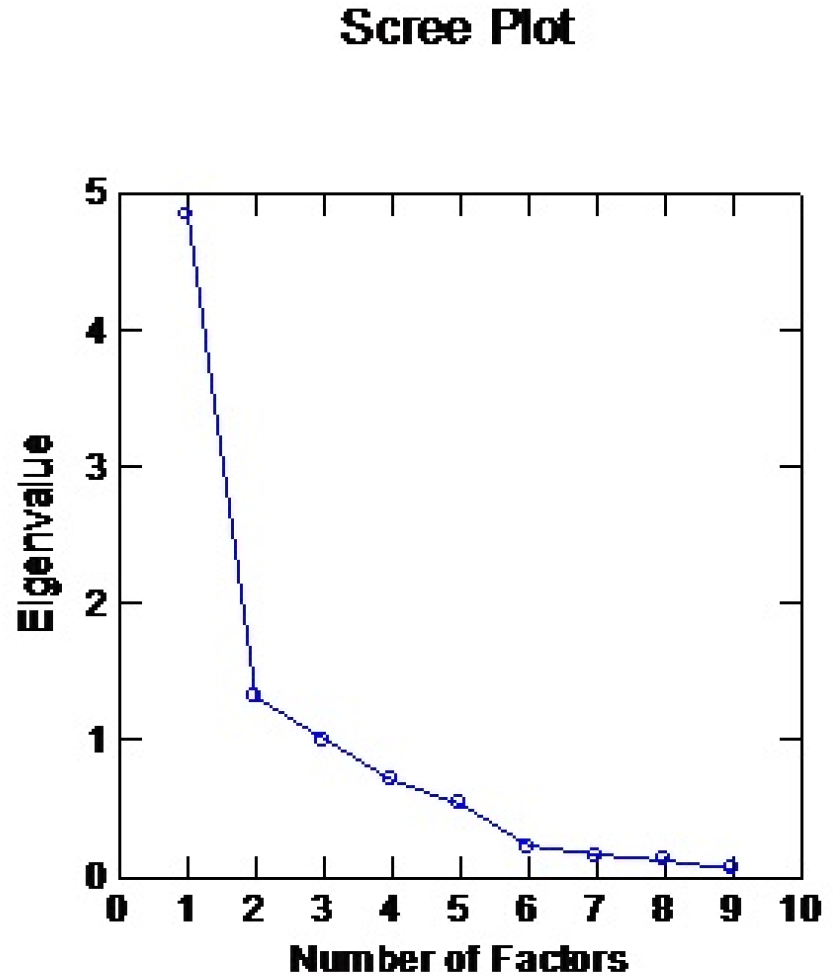
Scree plot obtained from factor analysis

**Fig 6b:**
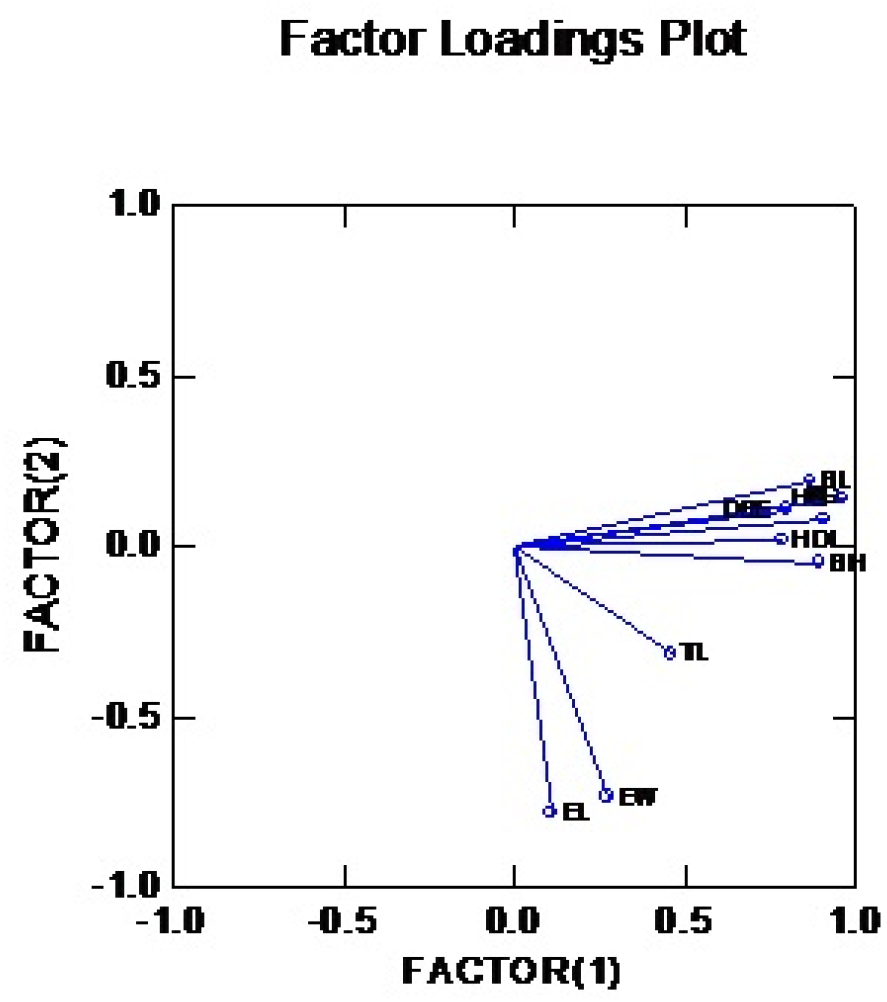
Factors loadings plot obtained from factor analysis

### Phylogenetic analysis of sheep reared under different agroclimatic regions of West Bengal by Hierarchical classification of cluster analysis

Hierarchical cluster analysis based on complete linkage, determine the Euclidean distance. The sheep population whose Euclidean distance was more, were clustered separately. The homogenous group of sheep population clustered together was observed to belong to a particular breed of sheep. Biodiversity study for sheep has been accessed through phylogenetic analysis of different sheep breeds of West Bengal along with their counterparts existing in eastern India (Fig. 7a). Different breeds of sheep usually found in Eastern India includes Garole, Chotanagpuri, Bonpala and Birbhum sheep of West Bengal; and Ganjam, Balangir, Sahabadi and Tibetan from other parts of the region. In the phylogenetic tree, different clusters depicts for different breeds of sheep as sequentially from top to bottom as Birbhum sheep, Garole, Chotanagpuri, Bonpala, Tibetan, Ganjam, Balangir, Kendrapada, and Sahabadi. It is evident from the constructed phylogenetic tree that the sheep of Hilly region as Tibetan and Bonpala were genetically distant from other sheep breeds of eastern India. Thus Bonpala is genetically most distant to other natives of West Bengal (i.e. Garole, Birbhum and Chotanagpuri). Phylogenetic analysis also reveals the possibility of slight genetic admixture of Garole and Chotanagpuri breeds which emphasizes the need for adoption of immediate conservation strategies for these two breeds of sheep to prevent the dilution of genetic merits. Garole and Birhum sheep found to exhibit much genetical closeness. Hence we had further analyzed these two breeds (Garole and Birbhum) covering a large number of populations (Fig. 7b), where they found to form two distinct clusters. It strengthens our hypothesis of genetic uniqueness of Garole and Birbhum sheep. Since the sheep existing in Birbhum district (dry-arid region) form a separate cluster and seems to form the basis for emerging a new breed of sheep. Ganjam, Kendrapada and Balangir (of Orissa) depicts genetic closeness. Sahabadi sheep were clustered separately. Genetic distance and detail of clusters by Hierarchical classification have been presented in *Supplementary Table 1*.

**Fig 7a:**
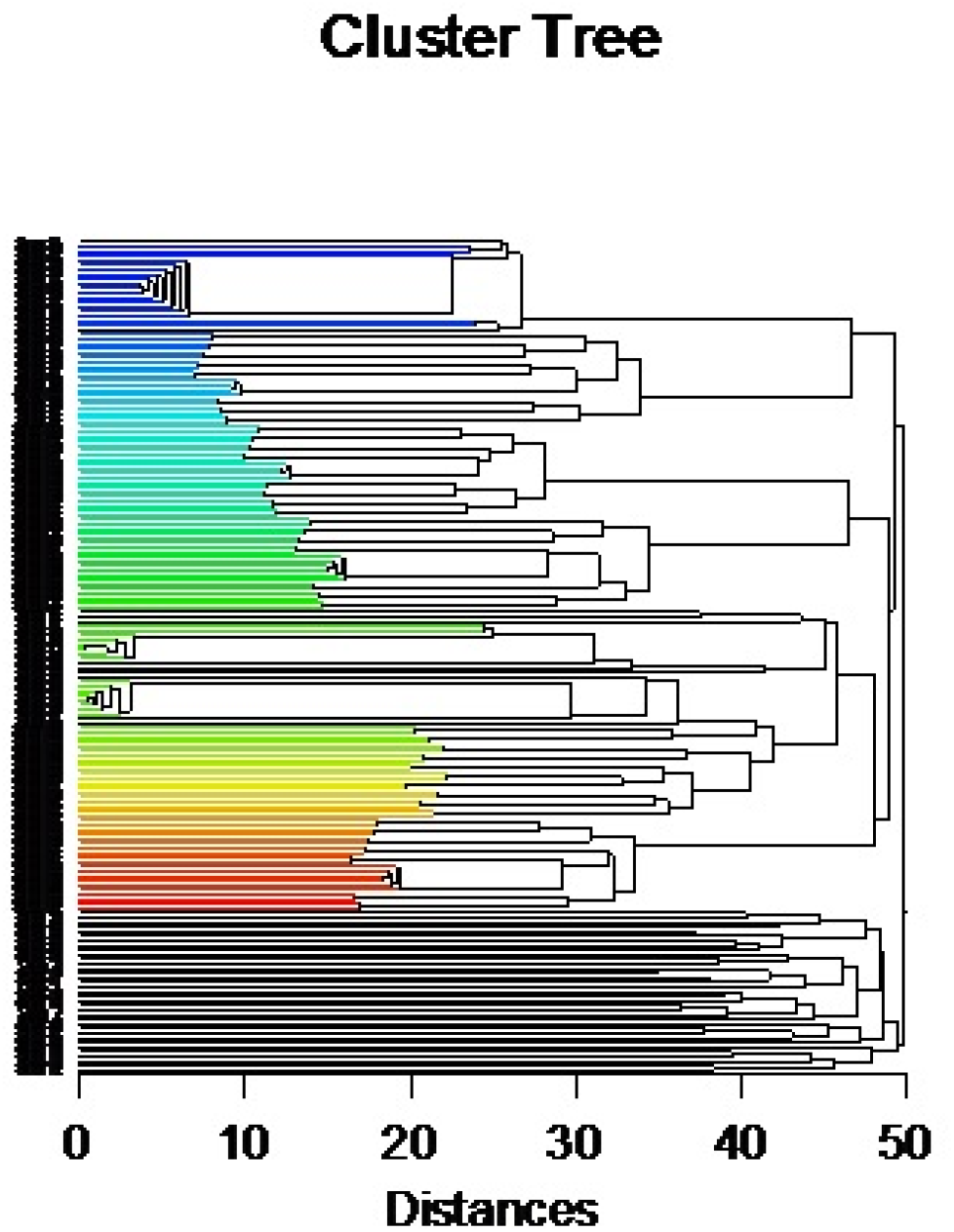
Phylogenetic tree obtained from sheep breeds from different agroclimatic conditions. **Phylog enetic tree for different sheep breeds of Eastern India (*Cluster Analysis-Hierarchial classification)*** Clusters in top to Bottom: 1. Blue: Birbhum 2. Light blue: Garole 3. Green: Chotanagpuri 4. Bonpala 5. Tibetan 6. Balangir 7. Ganjam 8. Kendrapada 9. Sahabadi

**Fig 7b:**
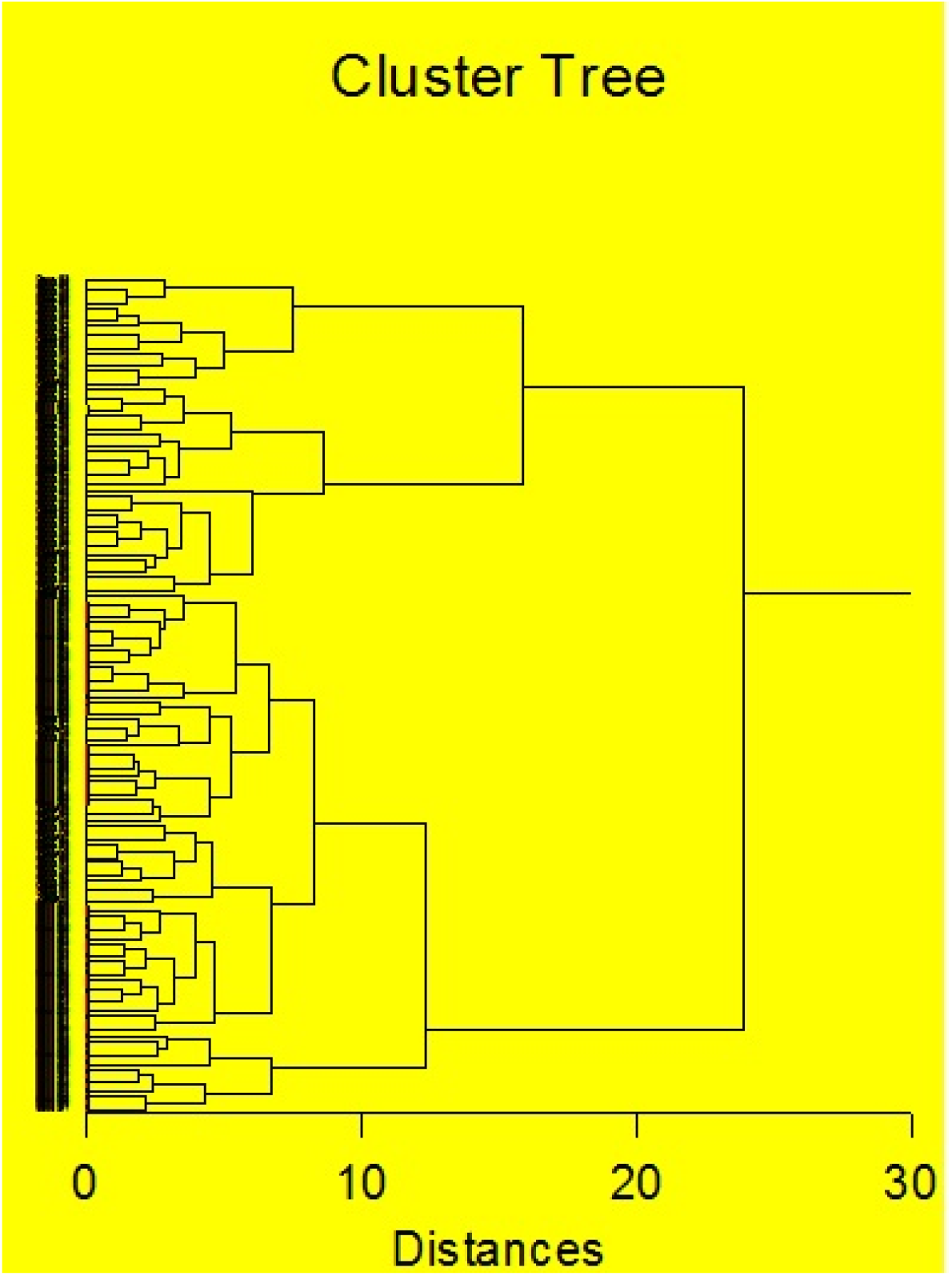
Phylogenetic tree for Garole and Birbhum sheep

### Phylogenetic analysis of sheep by K means cluster analysis

K means cluster analysis was employed for further differentiating Garole and Birbhum sheep based on biomorphometry. The figure 8 reveals two distinct clusters pertaining to two different sheep breeds as Birbhum and Garole based on identified biomorphometric characters as Heart girth, body length, body height and paunch girth. Table 8 represent the statistical analysis by K-means cluster based on identified biomorphometric traits. Genetic distance and detail of clusters by k-means. Cluster analysis have been presented in *Supplementary Table 2*.

**Fig 8:**
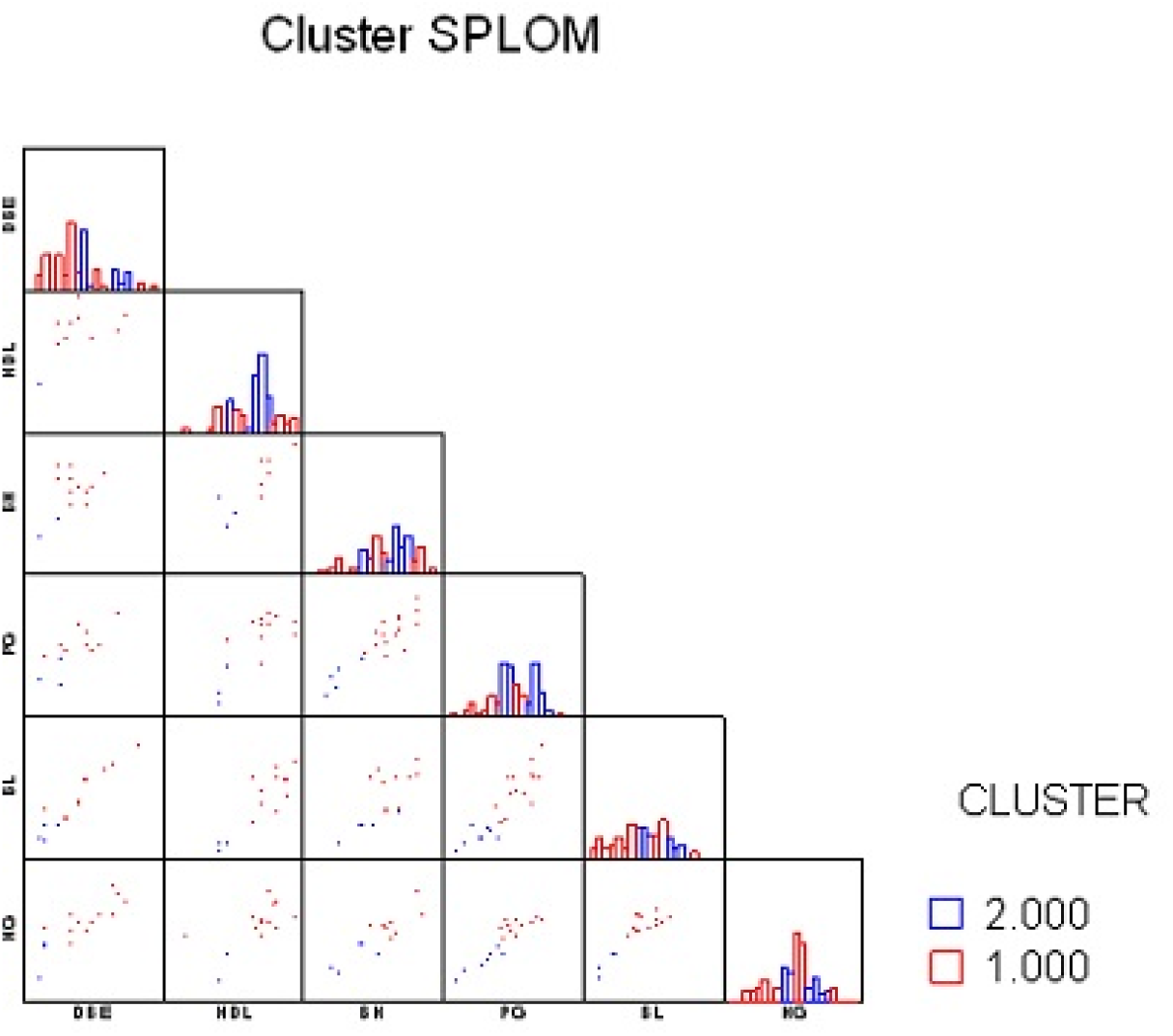
k-means cluster depicting different clusters for Garole and Birbhum

**Table 8:** K – means Cluster analysis for Birbhum and Garole sheep based on identified biomorphometric characters. K-Means Clustering **Distance Metric: Euclidean Distance** **K-Means Splitting Cases into 2 Groups**

**K-Means Splitting Cases into 2 Groups**
| Summary Statistics for All Cases |  |  |  |  |  |
| --- | --- | --- | --- | --- | --- |
| Variable | Between SS | df | Within SS | df | F-Ratio |
| HG | 5,044.191 | 1 | 3,204.351 | 91 | 143.249 |
| BL | 3,233.533 | 1 | 2,647.628 | 91 | 111.138 |
| BH | 1,540.994 | 1 | 1,618.188 | 91 | 86.659 |
| PG | 4,674.751 | 1 | 3,938.540 | 91 | 108.010 |
| HDL | 249.465 | 1 | 331.486 | 91 | 68.484 |
| DBE | 127.392 | 1 | 211.920 | 91 | 54.703 |
| ** TOTAL ** | 14,870.327 | 6 | 11,952.114 | 546 |  |

### Microsatellite genotyping for Ovis aries for Genetic Diversity Analysis

Measurements of within breed genetic variations as polymorphism information content (PIC) of different microsatellite loci and within population inbreeding estimates (F_IS_) are given in Table 5. The microsatellite loci amplified were observed to be polymorphic in the investigated Birbhum sheep population. All the markers were found to be highly informative with average PIC value of 0.693294. This indicates that these markers are quite useful for genetic diversity analysis. A total of 113 distinct alleles were identified across the 19 markers in Birbhum sheep. The observed no. of alleles ranged from 2 (BM1314, OarAE129) to 15 (CSSM31) with a mean of 6.27. The estimates of genetic diversity (mean expected heterozygosity) implied the presence of substantial amount of genetic variability in the Birbhum sheep population. The genetic variabilities in terms of microsatellite loci for Birbhum shhepis depicted in Table 9.

**Table 9:** Genetic variability measures among the Birbhum sheep based on microsatellite markers

| Marker | Major Allele Frquency | Genotype No | Allele No | Gene Diversity | PIC |
| --- | --- | --- | --- | --- | --- |
| BM6506 | 0.3333 | 3 | 4 | 0.7222 | 0.6713 |
| BM6526 | 0.25 | 4 | 6 | 0.8125 | 0.7861 |
| CSSM31 | 0.1111 | 8 | 15 | 0.9259 | 0.921 |
| BM1314 | 0.5 | 1 | 2 | 0.5 | 0.375 |
| BM757 | 0.3571 | 4 | 6 | 0.7245 | 0.6812 |
| INRA0063 | 0.1667 | 6 | 9 | 0.8704 | 0.8564 |
| MAF0214 | 0.4375 | 4 | 4 | 0.6484 | 0.5815 |
| OarFCB48 | 0.4 | 5 | 6 | 0.725 | 0.6833 |
| OarVH72 | 0.2778 | 5 | 7 | 0.8148 | 0.79 |
| OarHH64 | 0.5 | 3 | 4 | 0.6563 | 0.605 |
| OarJMP29 | 0.5 | 2 | 3 | 0.5938 | 0.5112 |
| OarCP20 | 0.2 | 10 | 10 | 0.87 | 0.8567 |
| oarJMP8 | 0.2778 | 6 | 8 | 0.8395 | 0.8211 |
| oarHH35 | 0.1667 | 8 | 12 | 0.8951 | 0.886 |
| oarCP0049 | 0.3 | 4 | 7 | 0.82 | 0.7978 |
| OarFCB128 | 0.3333 | 3 | 5 | 0.7639 | 0.726 |
| CSRD0247 | 0.5 | 2 | 3 | 0.625 | 0.5547 |
| OarAE129 | 0.5 | 1 | 2 | 0.5 | 0.375 |
| Mean |  |  |  | 0.739294 | 0.693294 |

Phylogenetic tree was constructed for the identified breeds from West Bengal. Chotanagpuri and Garole were clustered together. Bonpala and Birbhum were evolved as genetically distinct sheep breed (Fig 9).

**Fig 9:**
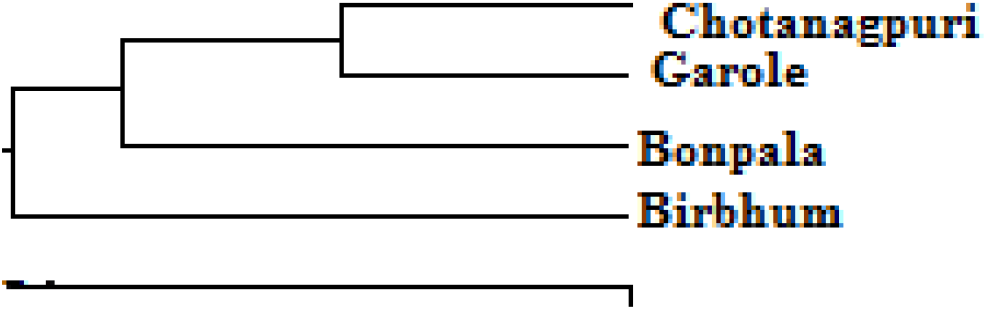
Phylogenetic tree constructed from Molecular evolutionary studies of Birbhum sheep w.r.t other sheep breeds of West Bengal

## 4. Discussion

Birbhum sheep has been identified as the newly reported sheep breed of West Bengal, which was reported to be clustered distinctly from other eight sheep breeds of eastern India (includes West Bengal, Bihar, Jharkhand and Odisha). In the present study 30.6 7% of the Birbhum sheep found to have rudimentary ear, which may be an unique characteristics for this particular breed. Body colour varied from black, brown and white for the Birbhum sheep. Horn is present in ram but absent in ewe in both Garole and Birbhum breed. Average ear length and ear width was exceptionally less for Birbhum in comparison to Garole might be due to higher percentage of rudimentary ear. Earlier a preliminary study was conducted in our lab^**2**^ and an idea was perceived in the similar way. Then in the current study we had analyzed more number of animals with a wide range of statistical techniques. Characterization of cattle breed based on growth and biomorphometric characteristics have been conducted for red Kandhari, Kankrej, Kenkatha, Red Sindhi ^**39-43**^.

Pearson correlation co-efficient reveals significant positive correlation for most of the growth traits in sheep at 12 months of age under study. Body weight was highly and positively correlated with biomorphometric traits as body length, body height, heart girth, punch girth and head length. Correlation between heart girth and body weight was found to be significantly higher at 12 months of age. Hence, at one year of age, in field condition, when weighing balance is not available, body weight can be easily accessed from heart girth, body length, paunch girth and body height.

Correlation of the growth traits has opened the avenue for PCA for the reduction of a set of observable variables. Phenotypic correlation among the different growth parameters including morphometric traits indicate similar high and positive correlation in dairy cattle and buffalo^**15**^. The PCA was applied to ten biomorphometric traits in sheep population of West Bengal from different agroclimatic zone of which, two components were extracted with that have eigen value greater than 1. Graphical representation with Scree plot depicts the various components having eigen values up to the point “bent of elbow” are usually considered. The identified two components could explain cumulative percentage of variance of 84.67%. First component accounted for 66.67% of the variation. It was represented by significant positive high loading of heart girth, body height, body length, punch girth, head length, distance between eyes and body weight. First component seemed to be explaining the maximum variability among the sheep. Our study showed better to that of <u>Salako (2006)</u>^44^ who also extracted two factors from 10 different biometric traits in Uda sheep which accounted for 75% of total variation. Principal component analysis of the morphostructural indices of White Fulani cattle had been conducted^**45-47**^

The observation was similar to Principal component analysis. Only two components have been identified with eigen value greater than one. Scree plot also depict the graphical representation and components having eigen values up to the point “bent of elbow” are usually considered. The identified two components could explain cumulative percentage of variance of 68.64%. First component accounted for 53.93% of the variation. It was represented by significant positive high loading of heart girth, body height, body length, punch girth, head length, distance between eyes and body weight. First component seemed to be explaining the maximum variability among the sheep. Factor analysis has been employed for sheep by a number of workers in different parts of the country^**48,49**^. Factor analysis of biometric traits of Kankrej cattle was studied^**42**^.

Hierarchical classification is based on Euclidean distance estimation (an indicator for genetic distance) for the sheep population under study. The individuals with lesser Euclidean distance were clustered together, indicating genetic similarity. All the individuals within a breed were presumed to be genetically similar. Individuals with genetically related breeds were clustered together. Biodiversity exists among different breeds of sheep usually found in Eastern India includes Garole, Chotanagpuri, Bonpala and Birbhm sheep of West Bengal; and Ganjam, Balangir, Sahabadi and Tibetan from other parts of the region. It is evident from the constructed phylogenetic tree that the sheep of Hilly region as Tibetan and Bonpala were genetically distant from other sheep breeds of eastern India. Phylogenetic analysis also reveals the possibility of slight genetic admixture of Garole and Chotanagpuri breeds, which emphasizes the need for adoption of immediate conservation strategies for these two breeds of sheep to prevent the dilution of genetic merits. Garole and Birhum sheep found to exhibit much genetical closeness. Hence we had further analyzed these two breeds (Garole and Birbhum) covering a large number of populations, where they found to form two distinct clusters. It strengthens our hypothesis of genetic uniqueness of Garole and Birbhum sheep. Since the sheep existing in Birbhum district (dry-arid region) form a separate cluster and seems to form the basis for emerging a new breed of sheep. Ganjam, Kendrapada and Balangir (of Orissa) depict genetic closeness. Sahabadi sheep were clustered separately. Genetic distance and detail of clusters by Hierarchical classification have been presented in *Supplementary Table 1*. Multivariate analysis of morphological characteristics in Nigerian native sheep have been studied in recent past^**46,47**^, where different clusters were identified for breeds of sheep. Genetic diversity study for sheep has also been conducted using the similar tool in other countries like Ethiopia^**50**^, and other European and Middle-Eastern sheep breeds^**51**^. Sheep biodiversity from Southern peninsular and parts of Eastern regions of India had been studied by microsatellite DNA marker^**52**^.

K-means cluster analysis is an analytical tool used with unlabeled data (i.e., data without defined categories or groups)and find groups in the data, with the number of groups (K). Data points are assigned and clustered based on feature similarity. Two distinct clusters were identified pertaining to two different sheep breeds as Birbhum and Garole based on identified biomorphometric characters by K means cluster analysis. K-means cluster analysis have been useful for biodiversity studies among the sheep population in different parts of the World^**53,54**^. They have predicted nonhierarchical clustering methods (e.g k-Means) as more preferable than hierarchical clustering methods in cluster analysis of two sheep breeds as Karayaka and Bafra. K-means cluster analysis has been highly effective in analysis of intensive Chios dairy sheep farms in Greece^**54**^.

In the current study, we could depict the distinct breed identity for Birbhum sheep, a newly reported breed through microsatellite genotyping. The present study implies the presence of substantial amount of genetic variability in the Birbhum sheep population. Microsatellite has been a popular genomic tool to analyze phylogenetic analysis in sheep in India^**55,57**^ and globally^56,58,59^.

## Conclusion

Genetic diversity exists within the existing breeds of sheep in West Bengal and four genetically distinct sheep breeds-Garole, Bonpala, Birbhum and Chotanagpuri were confirmed. Sheep reared at dry arid region of the Birbhum district may emerge as a new breed. A high positive significant phenotypic correlation was observed for growth and biomorphometric traits. Principal component analysis for 10 biomorphometric traits revealed two principal components with PC1 as the most promising. Factor analysis has also confirmed the above facts. Phylogenetic analysis by cluster analysis (both by hierarchical and K-means) also reveals Birbhum sheep as a separate cluster.Phylogenomic analysis with microsatellite genotyping also confirms the above findings.

Birbhum sheep were originated in the Birbhum district of West Bengal and they are known for more than 100 years, reared mostly for meat purpose. Body colour ranges from white, grey, to deep brown, either intact or patchy. They have a slightly convex head, with horizontal ears, two horns (blackish-grey colored) present only in ram but not in ewe; and absence of wattles. The head, face, and legs were almost devoid of wool, whereas rest of the body remains covered by wool of medium staple length, non-lustrous, straight or low crimp. The wool obtained as a by-product may be utilized for carpet wool production. Beard is absent and the tail is drooping downwards. The peculiarity of the Birbhum sheep is that 30.6% of sheep had a rudimentary ear with high disease resistance ability and better litter size. Hence, the rudimentary ear may be one of the breed characteristics of these sheep.

## Acknowledgement

Necessary funding provided by Department of Biotechnology, Govt. of India vide grant No. BT/Bio-CARe/04/10100/2013-14 and the support provided by Vice-Chancellor, West Bengal University of Animal and Fishery Sciences, Kolkata-37 as well as the Director, Animal Resource Development Department, Government of West Bengal, India is gratefully acknowledged.

The authors declare no conflict of interest.

## Notes

### Competing Interest Statement

The authors have declared no competing interest.

