## Supplementary material for "Phylophenomic and Phylogenomic analysis for *Ovis aries* reveals distinct identity of newly reported breed": Supl.Table1,2

Supplementary Table1: Clusters for Hierarchial clustering

►Hierarchical Clustering (supplementary)

| Clusters Joining | | at Distance | No. of Members |
|---|---|---|---|
| Case 33 | Case 1 | 0.000 | 2 |
| Case 94 | Case 87 | 0.000 | 2 |
| Case 106 | Case 96 | 0.000 | 2 |
| Case 99 | Case 97 | 0.000 | 2 |
| Case 103 | Case 99 | 0.000 | 3 |
| Case 105 | Case 103 | 0.000 | 4 |
| Case 107 | Case 105 | 0.000 | 5 |
| Case 108 | Case 106 | 0.000 | 3 |
| Case 109 | Case 107 | 0.000 | 6 |
| Case 110 | Case 108 | 0.000 | 4 |
| Case 111 | Case 109 | 0.000 | 7 |
| Case 112 | Case 110 | 0.000 | 5 |
| Case 113 | Case 111 | 0.000 | 8 |
| Case 114 | Case 112 | 0.000 | 6 |
| Case 115 | Case 113 | 0.000 | 9 |
| Case 116 | Case 114 | 0.000 | 7 |
| Case 117 | Case 115 | 0.000 | 10 |
| Case 118 | Case 116 | 0.000 | 8 |
| Case 119 | Case 117 | 0.000 | 11 |
| Case 120 | Case 118 | 0.000 | 9 |
| Case 121 | Case 119 | 0.000 | 12 |
| Case 122 | Case 120 | 0.000 | 10 |
| Case 123 | Case 121 | 0.000 | 13 |
| Case 124 | Case 122 | 0.000 | 11 |
| Case 125 | Case 123 | 0.000 | 14 |
| Case 126 | Case 124 | 0.000 | 12 |
| Case 127 | Case 125 | 0.000 | 15 |
| Case 128 | Case 126 | 0.000 | 13 |
| Case 129 | Case 127 | 0.000 | 16 |
| Case 130 | Case 128 | 0.000 | 14 |
| Case 131 | Case 129 | 0.000 | 17 |
| Case 132 | Case 130 | 0.000 | 15 |
| Case 133 | Case 131 | 0.000 | 18 |
| Case 134 | Case 132 | 0.000 | 16 |
| Case 135 | Case 133 | 0.000 | 19 |
| Case 136 | Case 134 | 0.000 | 17 |
| Case 137 | Case 135 | 0.000 | 20 |
| Case 138 | Case 136 | 0.000 | 18 |
| Case 139 | Case 137 | 0.000 | 21 |
| Case 140 | Case 138 | 0.000 | 19 |
| Case 141 | Case 139 | 0.000 | 22 |
| Case 142 | Case 140 | 0.000 | 20 |
| Case 143 | Case 141 | 0.000 | 23 |

| Clusters Joining | | at Distance | No. of Members |
|---|---|---|---|
| Case 144 | Case 142 | 0.000 | 21 |
| Case 145 | Case 143 | 0.000 | 24 |
| Case 147 | Case 146 | 0.000 | 2 |
| Case 148 | Case 147 | 0.000 | 3 |
| Case 149 | Case 148 | 0.000 | 4 |
| Case 150 | Case 149 | 0.000 | 5 |
| Case 151 | Case 150 | 0.000 | 6 |
| Case 152 | Case 151 | 0.000 | 7 |
| Case 153 | Case 152 | 0.000 | 8 |
| Case 154 | Case 153 | 0.000 | 9 |
| Case 155 | Case 154 | 0.000 | 10 |
| Case 156 | Case 155 | 0.000 | 11 |
| Case 157 | Case 156 | 0.000 | 12 |
| Case 158 | Case 157 | 0.000 | 13 |
| Case 159 | Case 158 | 0.000 | 14 |
| Case 160 | Case 159 | 0.000 | 15 |
| Case 144 | Case 100 | 0.006 | 22 |
| Case 144 | Case 98 | 0.017 | 23 |
| Case 145 | Case 95 | 0.098 | 25 |
| Case 102 | Case 144 | 0.156 | 24 |
| Case 104 | Case 102 | 0.393 | 25 |
| Case 101 | Case 145 | 0.512 | 26 |
| Case 71 | Case 65 | 0.577 | 2 |
| Case 79 | Case 54 | 0.645 | 2 |
| Case 24 | Case 2 | 0.816 | 2 |
| Case 17 | Case 6 | 0.816 | 2 |
| Case 42 | Case 23 | 0.816 | 2 |
| Case 31 | Case 29 | 0.816 | 2 |
| Case 43 | Case 38 | 0.816 | 2 |
| Case 89 | Case 76 | 0.816 | 2 |
| Case 92 | Case 78 | 0.816 | 2 |
| Case 37 | Case 28 | 1.000 | 2 |
| Case 73 | Case 48 | 1.000 | 2 |
| Case 70 | Case 69 | 1.000 | 2 |
| Case 62 | Case 101 | 1.050 | 27 |
| Case 12 | Case 24 | 1.155 | 3 |
| Case 39 | Case 21 | 1.155 | 2 |
| Case 88 | Case 73 | 1.155 | 3 |
| Case 46 | Case 4 | 1.291 | 2 |
| Case 32 | Case 22 | 1.291 | 2 |
| Case 83 | Case 34 | 1.291 | 2 |
| Case 49 | Case 41 | 1.291 | 2 |
| Case 57 | Case 56 | 1.291 | 2 |
| Case 82 | Case 67 | 1.291 | 2 |
| Case 86 | Case 85 | 1.323 | 2 |
| Case 47 | Case 5 | 1.414 | 2 |
| Case 11 | Case 7 | 1.414 | 2 |
| Case 35 | Case 16 | 1.414 | 2 |

| Clusters Joining | | at Distance | No. of Members |
|---|---|---|---|
| Case 42 | Case 40 | 1.414 | 3 |
| Case 70 | Case 52 | 1.414 | 3 |
| Case 94 | Case 92 | 1.414 | 4 |
| Case 14 | Case 13 | 1.425 | 2 |
| Case 19 | Case 8 | 1.633 | 2 |
| Case 36 | Case 26 | 1.633 | 2 |
| Case 45 | Case 44 | 1.633 | 2 |
| Case 79 | Case 66 | 1.633 | 3 |
| Case 14 | Case 37 | 1.662 | 4 |
| Case 33 | Case 17 | 1.732 | 4 |
| Case 64 | Case 71 | 1.732 | 3 |
| Case 93 | Case 68 | 1.732 | 2 |
| Case 30 | Case 3 | 1.826 | 2 |
| Case 88 | Case 72 | 1.826 | 4 |
| Case 20 | Case 19 | 1.915 | 3 |
| Case 53 | Case 51 | 1.915 | 2 |
| Case 84 | Case 82 | 2.000 | 3 |
| Case 35 | Case 15 | 2.082 | 3 |
| Case 47 | Case 18 | 2.160 | 3 |
| Case 42 | Case 49 | 2.160 | 5 |
| Case 77 | Case 55 | 2.160 | 2 |
| Case 64 | Case 63 | 2.160 | 4 |
| Case 25 | Case 10 | 2.309 | 2 |
| Case 59 | Case 104 | 2.359 | 26 |
| Case 32 | Case 12 | 2.380 | 5 |
| Case 83 | Case 39 | 2.380 | 4 |
| Case 91 | Case 80 | 2.380 | 2 |
| Case 79 | Case 94 | 2.398 | 7 |
| Case 74 | Case 77 | 2.449 | 3 |
| Case 27 | Case 9 | 2.582 | 2 |
| Case 60 | Case 58 | 2.646 | 2 |
| Case 32 | Case 30 | 2.944 | 7 |
| Case 90 | Case 84 | 3.000 | 4 |
| Case 59 | Case 62 | 3.109 | 53 |
| Case 33 | Case 11 | 3.162 | 6 |
| Case 31 | Case 27 | 3.162 | 4 |
| Case 88 | Case 86 | 3.227 | 6 |
| Case 25 | Case 43 | 3.317 | 4 |
| Case 50 | Case 57 | 3.317 | 3 |
| Case 70 | Case 64 | 3.317 | 7 |
| Case 74 | Case 89 | 3.464 | 5 |
| Case 32 | Case 42 | 3.916 | 12 |
| Case 60 | Case 53 | 4.000 | 4 |
| Case 50 | Case 59 | 4.022 | 56 |
| Case 14 | Case 36 | 4.062 | 6 |
| Case 20 | Case 46 | 4.123 | 5 |
| Case 93 | Case 47 | 4.203 | 5 |
| Case 81 | Case 61 | 4.243 | 2 |

| Clusters Joining | | at Distance | No. of Members |
|---|---|---|---|
| Case 45 | Case 25 | 4.359 | 6 |
| Case 74 | Case 79 | 4.435 | 12 |
| Case 70 | Case 91 | 4.509 | 9 |
| Case 31 | Case 35 | 4.761 | 7 |
| Case 20 | Case 32 | 5.477 | 17 |
| Case 33 | Case 93 | 5.508 | 11 |
| Case 90 | Case 88 | 5.575 | 10 |
| Case 83 | Case 45 | 6.164 | 10 |
| Case 60 | Case 50 | 7.239 | 60 |
| Case 90 | Case 74 | 7.528 | 22 |
| Case 20 | Case 33 | 8.103 | 28 |
| Case 31 | Case 14 | 8.505 | 13 |
| Case 81 | Case 70 | 8.737 | 11 |
| Case 90 | Case 20 | 10.283 | 50 |
| Case 81 | Case 90 | 12.910 | 61 |
| Case 31 | Case 83 | 15.351 | 23 |
| Case 81 | Case 60 | 19.225 | 121 |
| Case 81 | Case 160 | 34.109 | 136 |
| Case 31 | Case 81 | 49.070 | 159 |

Table 2: Different clusters by K means. (supplementary file)

| Cluster 1 of 2 Contains 73 Cases | | | | | | |
|---|---|---|---|---|---|---|
| Members | | Statistics | | | | |
| Case | Distance | Variable | Minimum | Mean | Maximum | Standard Deviation |
| Case 1 | 3.723 | HG | 48.000 | 58.315 | 74.000 | 6.105 |
| Case 2 | 3.381 | BL | 29.000 | 38.877 | 53.000 | 5.845 |
| Case 3 | 4.777 | BH | 39.000 | 46.658 | 56.000 | 4.022 |
| Case 4 | 4.366 | PG | 47.000 | 58.856 | 76.000 | 6.570 |
| Case 5 | 2.805 | HDL | 7.000 | 15.986 | 20.000 | 2.039 |
| Case 6 | 3.638 | DBE | 6.000 | 9.199 | 14.000 | 1.670 |
| Case 7 | 3.881 | | | | | |
| Case 8 | 3.252 | | | | | |
| Case 11 | 3.291 | | | | | |
| Case 12 | 3.583 | | | | | |
| Case 17 | 4.229 | | | | | |
| Case 18 | 1.628 | | | | | |
| Case 19 | 2.988 | | | | | |
| Case 20 | 3.310 | | | | | |
| Case 21 | 6.504 | | | | | |
| Case 22 | 3.314 | | | | | |

| Cluster 1 of 2 Contains 73 Cases | | | | | | |
|---|---|---|---|---|---|---|
| Members | | Statistics | | | | |
| Case | Distance | Variable | Minimum | Mean | Maximum | Standard Deviation |
| Case 23 | 4.240 | | | | | |
| Case 24 | 3.963 | | | | | |
| Case 30 | 2.832 | | | | | |
| Case 32 | 3.739 | | | | | |
| Case 33 | 2.762 | | | | | |
| Case 34 | 6.103 | | | | | |
| Case 39 | 6.429 | | | | | |
| Case 40 | 5.932 | | | | | |
| Case 41 | 4.847 | | | | | |
| Case 46 | 5.937 | | | | | |
| Case 47 | 2.952 | | | | | |
| Case 48 | 6.442 | | | | | |
| Case 49 | 3.617 | | | | | |
| Case 50 | 9.319 | | | | | |
| Case 51 | 9.220 | | | | | |
| Case 52 | 4.214 | | | | | |
| Case 53 | 7.079 | | | | | |
| Case 54 | 3.863 | | | | | |
| Case 55 | 5.126 | | | | | |
| Case 56 | 6.959 | | | | | |
| Case 57 | 9.031 | | | | | |
| Case 58 | 6.634 | | | | | |
| Case 59 | 8.594 | | | | | |
| Case 60 | 5.007 | | | | | |
| Case 61 | 5.849 | | | | | |
| Case 62 | 9.713 | | | | | |
| Case 63 | 4.233 | | | | | |
| Case 64 | 3.979 | | | | | |
| Case 65 | 4.285 | | | | | |
| Case 66 | 2.534 | | | | | |
| Case 67 | 3.738 | | | | | |
| Case 68 | 2.571 | | | | | |
| Case 69 | 3.418 | | | | | |
| Case 70 | 4.448 | | | | | |
| Case 71 | 2.804 | | | | | |
| Case 72 | 5.054 | | | | | |
| Case 73 | 4.218 | | | | | |
| Case 74 | 4.002 | | | | | |
| Case 75 | 2.143 | | | | | |
| Case 76 | 3.410 | | | | | |
| Case 77 | 2.114 | | | | | |
| Case 78 | 3.059 | | | | | |
| Case 79 | 4.294 | | | | | |
| Case 80 | 3.139 | | | | | |
| Case 81 | 3.305 | | | | | |
| Case 82 | 5.477 | | | | | |

| Cluster 1 of 2 Contains 73 Cases | | | | | | |
| --- | --- | --- | --- | --- | --- | --- |
| Members | | Statistics | | | | |
| Case | Distance | Variable | Minimum | Mean | Maximum | Standard Deviation |
| Case 83 | 2.612 | | | | | |
| Case 84 | 5.255 | | | | | |
| Case 85 | 5.121 | | | | | |
| Case 86 | 1.794 | | | | | |
| Case 87 | 5.482 | | | | | |
| Case 88 | 1.884 | | | | | |
| Case 89 | 3.870 | | | | | |
| Case 90 | 3.776 | | | | | |
| Case 91 | 3.078 | | | | | |
| Case 92 | 3.664 | | | | | |
| Case 93 | 3.621 | | | | | |

| Cluster 2 of 2 Contains 20 Cases | | | | | | |
| --- | --- | --- | --- | --- | --- | --- |
| Members | | Statistics | | | | |
| Case | Distance | Variable | Minimum | Mean | Maximum | Standard Deviation |
| Case 9 | 1.600 | HG | 32.000 | 40.390 | 51.000 | 5.237 |
| Case 10 | 4.566 | BL | 20.000 | 24.525 | 32.000 | 3.143 |
| Case 13 | 4.077 | BH | 29.000 | 36.750 | 48.000 | 4.887 |
| Case 14 | 2.576 | PG | 29.000 | 41.600 | 54.000 | 6.613 |
| Case 15 | 2.522 | HDL | 10.000 | 12.000 | 15.000 | 1.298 |
| Case 16 | 1.541 | DBE | 5.500 | 6.350 | 8.000 | 0.763 |
| Case 25 | 4.776 | | | | | |
| Case 26 | 5.601 | | | | | |
| Case 27 | 2.107 | | | | | |
| Case 28 | 3.424 | | | | | |
| Case 29 | 2.052 | | | | | |
| Case 31 | 3.217 | | | | | |
| Case 35 | 1.747 | | | | | |
| Case 36 | 7.197 | | | | | |
| Case 37 | 4.374 | | | | | |
| Case 38 | 2.901 | | | | | |
| Case 42 | 7.154 | | | | | |
| Case 43 | 3.375 | | | | | |
| Case 44 | 6.190 | | | | | |
| Case 45 | 3.977 | | | | | |
